# Structure-inspired design of Nsp8-based protein inhibitors to suppress SARS-CoV-2 replication

**DOI:** 10.64898/2026.08.30.747997

**Authors:** Petra Štěrbová, Hsin-Hung Lin, I-Ping Tu, Wei-Hau Chang

**Affiliations:** Institute of Chemistry, Academia Sinica, Taipei, 115, Taiwan; Institute of Statistical Science, Academia Sinica, Taipei, 115, Taiwan

## Abstract

SARS-CoV-2 relies on a conserved RNA-dependent RNA polymerase (RdRp) complex composed of nsp12 and its cofactors nsp7 and nsp8 to replicate its RNA genome. Whereas most antiviral strategies target viral enzymes or surface proteins directly, an alternative approach is to disrupt the assembly or function of an essential viral molecular machine using a defective component derived from the pathogen itself. Here, guided by structural analyses of the nsp12–nsp7–nsp8 replication complex, we designed truncated nsp8 proteins that retain the ability to associate with nsp12 but are defective in engaging RNA. Using a purified nsp12–nsp7–nsp8 system capable of RNA primer extension, we show that selected truncated nsp8 variants inhibit polymerase activity when introduced into an otherwise functional complex. These results are consistent with a competitive mechanism in which the defective nsp8 variants associate with nsp12 and interfere with incorporation or function of wild-type nsp8, thereby compromising formation of a productive replication complex. To further explore this strategy, we used structure-guided in silico analysis of the nsp8– nsp12 interface to identify interaction hotspots and screened corresponding single-amino-acid substitutions. Several variants exhibited enhanced inhibitory activity in the reconstituted polymerase assay. Together, these findings establish a proof-of-concept strategy in which a structurally engineered, pathogen-derived protein can act as a dominant-negative inhibitor of an essential viral replication machinery. This approach provides a framework for developing protein- or peptide-based inhibitors that target conserved protein–protein interactions within viral replication complexes.

## Introduction

The coronavirus disease 2019 (COVID-19) was caused by a global outbreak of a severe acute respiratory syndrome coronavirus 2 (SARS-CoV-2) [1]. SARS-CoV-2 belongs to the genus Betacoronavirus of the family *Coronaviridae*. Coronaviruses are enveloped viruses with a large genome consisting of a non-segmented positive sense, single stranded RNA [2].

The continued evolution of SARS-CoV-2 presents a persistent challenge for antiviral development. Viral determinants exposed to immune pressure can accumulate sequence variation, motivating strategies that target molecular functions constrained by essential steps of viral replication. The viral RNA-dependent RNA polymerase (RdRp) is particularly attractive in this regard because genome replication depends on its activity and on the assembly of a defined multiprotein complex [3]. This essential role of RdRp for virus replication makes viral polymerase a highly promising target for antiviral therapeutics [4]. Currently, amongst therapeutics approved for clinical usage against SARS-CoV-2 targeting RdRp catalytic site are nucleoside analogues Remdesivir and Molnupiravir [5-7]. The nucleoside analogues mimic the structure of natural RdRp substrates. However, the use of nucleoside analogues for treatment of COVID-19 patients have variable outcomes in efficacy and specificity [8,9]. Accordingly, in addition to targeting the catalytic activity of the polymerase, interfering with essential protein–protein interactions within the replication machinery may provide an alternative route for suppressing viral RNA synthesis.

The SARS-CoV-2 RdRp complex consists of the catalytic component of Nsp12 and the accessory factors Nsp7 and Nsp8. Nsp12 contains the conserved active site responsible for RNA synthesis, which is targeted by antiviral nucleoside analogs such as remdesivir and molnupiravir. However, productive RNA synthesis also depends on the structural organization and stabilization of Nsp12 by its accessory factors. Structural studies have revealed extensive interfaces between Nsp12, Nsp7, and Nsp8, with Nsp8 making extensive contacts with the polymerase and contributing to the architecture of the RNA-synthesis complex [10,11]. These observations suggest that the assembly interfaces of the replication machinery, in addition to the catalytic pocket, represent functionally constrained sites that may be susceptible to competitive inhibition.

The Nsp12–Nsp8 interface is particularly attractive for such an approach because an inhibitor occupying the Nsp8-binding site on Nsp12 could, in principle, compete directly with the native Nsp8 cofactor and thereby interfere with assembly of a functional polymerase complex. Analysis of the interactions between RdRp subunits revealed that the nsp12-nsp8 interaction possess the highest total interaction energy [12]. The interaction between nsp12 and nsp8 has been demonstrated to be associated with RdRp activity. SARS-CoV-2 variants containing the nsp12 mutation P323L exhibits enhanced RdRp enzymatic activity associated with increased RdRp complex stability and enzymatic activity [13]. Further mutations of nsp12 altering the interactions between nsp12-nsp8 interface resulted in impaired RdRp activity [14]. Therefore, a peptide-based or peptide-mimetic inhibitor capable of competitively occupying the nsp8 docking region on nsp12 could, in principle, dismantle the polymerase complex and suppress viral replication through a mechanism distinct from that of conventional active-site inhibitors.

Recent work has provided an important proof of principle for this concept: short synthetic peptides derived from the Nsp8–Nsp12 interface were shown to bind Nsp12 and inhibit SARS-CoV-2 polymerase activity [15]. These studies establish that an Nsp8-binding surface on Nsp12 can be exploited by molecular mimics of the native cofactor. However, they also leave open an important protein-engineering question. Rather than restricting the inhibitor to a short interface peptide, can a larger Nsp8-derived protein scaffold retain competitive activity while providing additional structural context that can be systematically engineered to improve inhibition?

We addressed this question using an Nsp8 fragment encompassing residues 76–198. This region retains the Nsp8 structural framework involved in interaction with Nsp12 while lacking the extended RNA-binding arm of the native protein. We reasoned that such a fragment could function as a recombinant competitive inhibitor by engaging the Nsp12 surface normally occupied by Nsp8 without contributing the RNA-binding activity required of the intact cofactor. This strategy differs from the short synthetic peptides described previously [15] and provides a protein-based scaffold in which individual residues at the Nsp12-binding interface can be experimentally modified.

A central challenge in engineering such an inhibitor is identifying substitutions that alter its competitive behavior without disrupting the structural integrity of the fragment. Structure prediction provides a means of systematically exploring this sequence space. In particular, AlphaFold3 can generate structural models of protein– protein complexes [16] and can therefore be used to evaluate how individual substitutions may influence the interaction of an Nsp8-derived fragment with Nsp12.

We used this capability as a structure-guided screening strategy to prioritize single-residue variants at the Nsp8–Nsp12 interface. Importantly, rather than assuming that computational scores directly report inhibitory potency, we used the predictions to generate experimentally testable hypotheses about sequence-dependent competition [17,18] with the native Nsp8 interaction.

Here, we experimentally evaluate Nsp8(76–198) and structure-guided single-residue variants using an in vitro RNA primer-extension assay that directly reports inhibition of the Nsp12–Nsp7–Nsp8 replication machinery. Most variants selected for favorable computational interaction scores showed inhibition profiles comparable to the parental fragment. Unexpectedly, however, a substitution at Nsp8 residue 104, despite only a modest computational score, produced markedly enhanced and saturable inhibition at approximately equimolar inhibitor-to-polymerase-complex stoichiometry. These findings reveal that computationally favorable interaction scores do not simply translate into stronger inhibition, while demonstrating that individual residues within the Nsp8–Nsp12 interface can profoundly influence the competitive behavior of an Nsp8-derived inhibitor. Our work therefore establishes a recombinant Nsp8-fragment framework for interrogating and engineering competitive inhibition of an essential protein–protein interaction within the SARS-CoV-2 replication machinery.

## Results

### *In vitro* reconstituted SARS-CoV-2 core RdRp elongates RNA substrate

The SARS-CoV-2 minimal replication machinery is composed of nsp12, the RdRp, nsp7, and two copies of nsp8 [10,19]. The nsp7 and nsp8 are cofactors to enable the RdRp for engaging long RNA substrate and stabilize the RdRp complex to enhance procressivity. To obtain nsp12, nsp7, and nsp8, we used *E. coli* to produce over-expressed proteins for purification (**Fig. 1A**). A RdRp complex was reconstituted *in vitro* by co-incubating nsp12, nsp7, and nsp8 at a 1:3:6 ratio. The activity of reconstituted RdRp was assessed using a fluorescence-based RNA extension assay with a RNA hairpin scaffold with Cy3 (Cyanine3) label at 5’ end (**Fig. 1B**). The *in vitro* elongation activity showed that individually produced subunits can be assembled into a functional RdRp complex to support RNA primer elongation (**Fig. 1C**).

**Figure 1.**
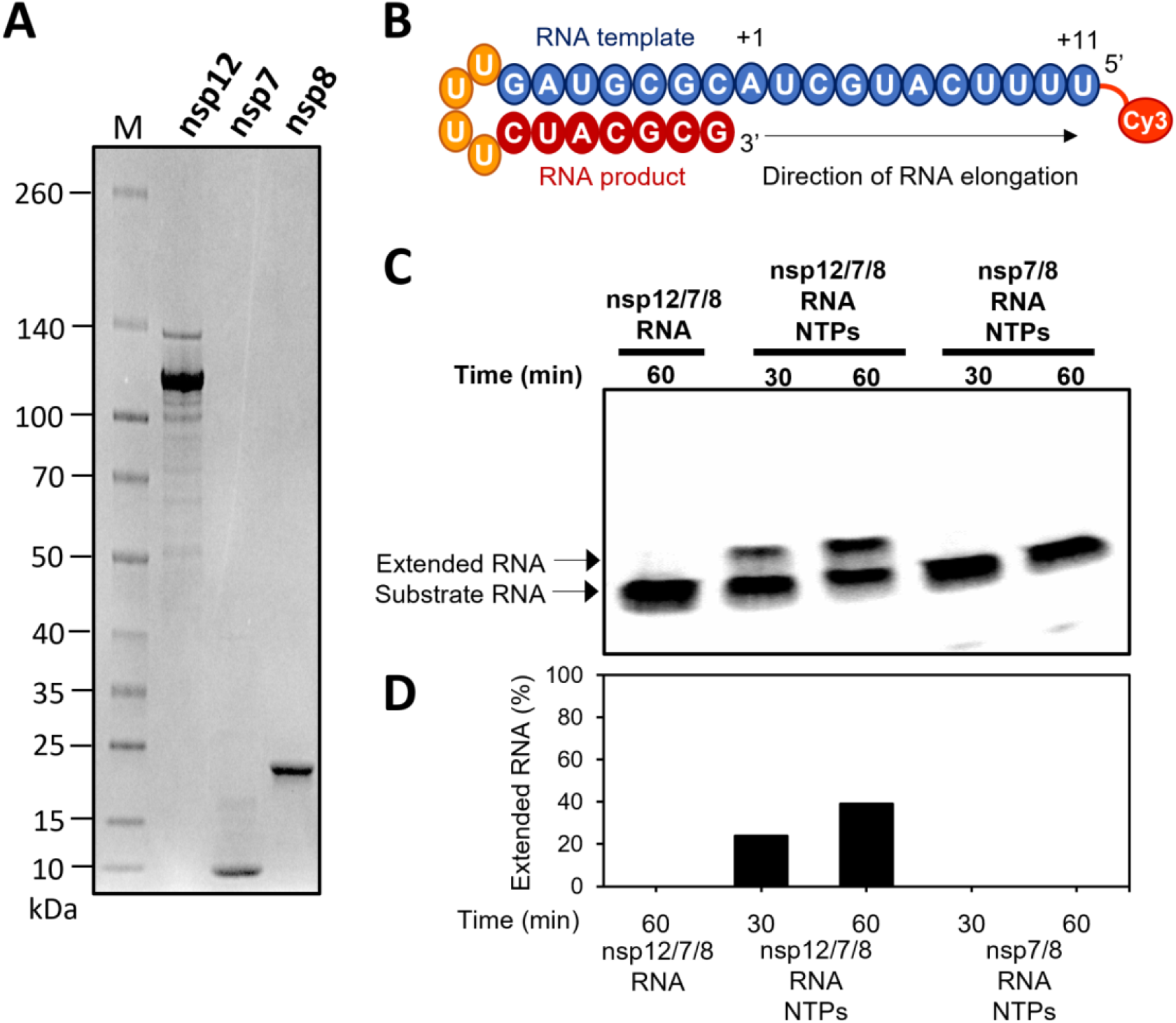
Reconstituted RdRp complex catalyzes RNA elongation in vitro. **(A)** SDS-PAGE analysis of the purified SARS-CoV-2 RdRp subunits nsp12, nsp7, and nsp8. **(B)** Minimal RNA hairpin substrate with fluorescently labelled 5’ overhang used for *in vitro* RdRp activity assay. **(C)** The reconstituted RdRp complex shows polymerase activity *in vitro*.

### nsp8 N-terminal arm is essential for supporting RNA elongation

As shown in **Fig. 2B**, the nsp12 RdRp RNA elongation activity required nsp7 and nsp8 co-factors. Mechanistic insights into this molecular requirement is revealed by the reported architecture of a replicating RdRp complex, in which a nsp7 and two copies of nsp8 associate with a nsp12 via protein-protein interactions, with nsp8’s arm-like structure extended to engage the RNA template (**Fig. 2A**). The nsp8 consists of two domains; an extension domain at the N-terminal consisting of a long α-helix forming protruding positively charged arm to engage the exiting RNA, and a head domain at the C-terminal that, in the first copy, interacts with nsp7, and in the second copy, interacts with nsp12 [10].

**Figure 2.**
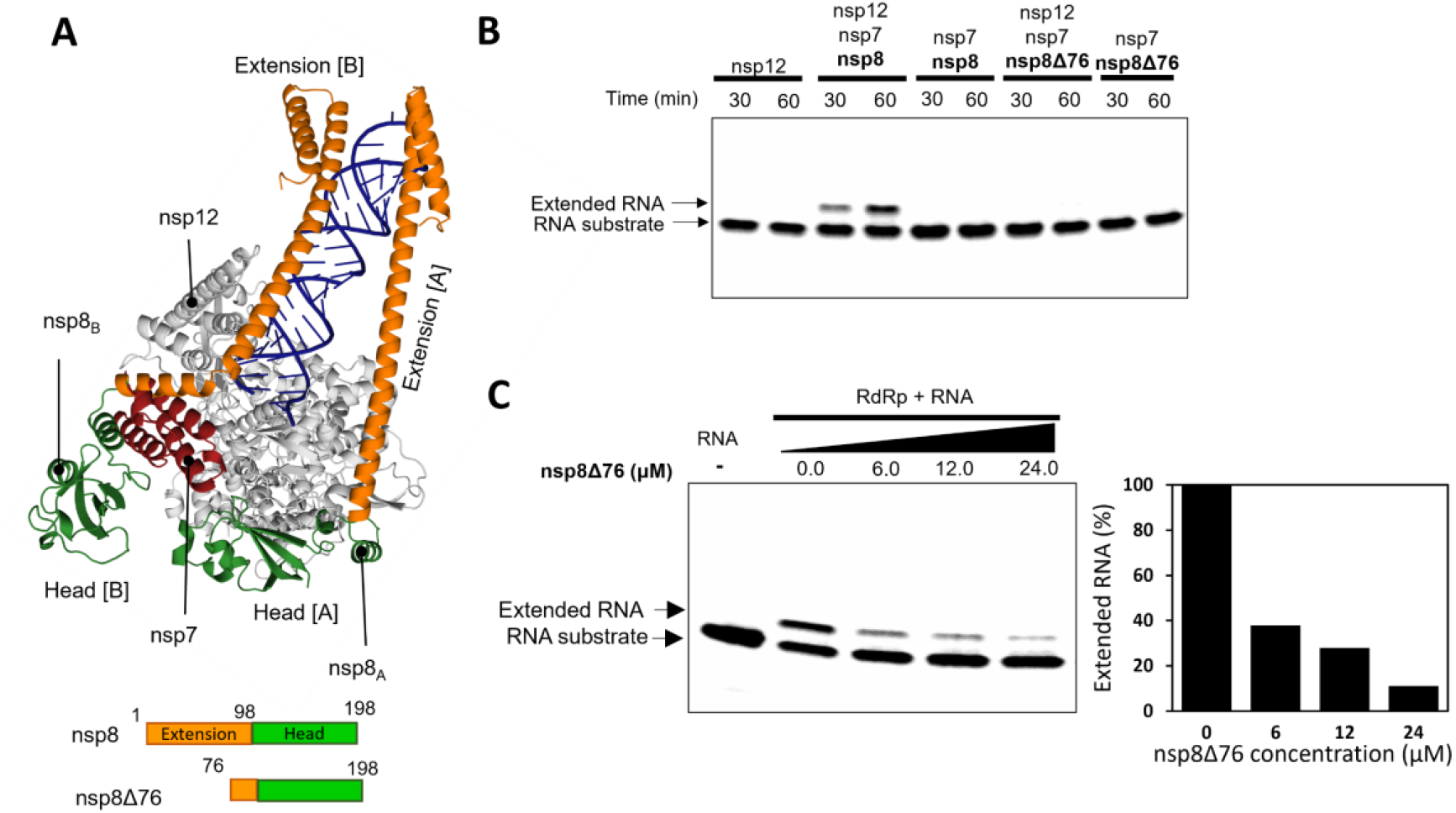
N-terminal α-helix of nsp8 is critical for RdRp activity. **(A)** Cartoon representation of SARS-CoV-2 core RdRp complex (PDB 6YYT). The nsp8 extension domain is coloured in orange and head domain in green. **(B)** nsp8Δ76 construct does not promote RNA elongation by RdRp. **(C)** Left: Addition of nsp8Δ76 to RNA elongation reaction inhibits the RNA product elongation. Right: Quantification of the RNA product.

We thus hypothesized that an nsp8 lacking N-terminal extensions, for example, an nsp8 mutant Δ1-75 (nsp8Δ76), would render the RdRp to lose RNA elongation capability. To test its idea, we created nsp8Δ76 that lacks the RNA interacting N-terminal domain, but retains the C-terminal domain that interacts with nsp7 or nsp12. As shown in **Figure 2B**, the RdRp complex reconstructed using this truncated nsp8 indeed failed to support RNA elongation activity.

To further test whether the truncated nsp8 has a potential to inhibit RNA elongation, we added this nsp8Δ76 to the minimal RdRp replicating system consisting of nsp12, nsp7 and nsp8 that support RNA elongation. The addition of nsp8Δ76 lead to a dose-dependent decrease of formation RNA elongation product (**Fig. 2C**). The inhibition in the presence of nsp8Δ76 seems to be quantitative—addition of 6 μM of the nsp8Δ76 (equal amount of the full length nsp8) resulted in approximately 2-fold reduction of the RNA elongation product while addition of 24 μM of the nsp8Δ76 (4-fold of the full length) resulted in approximately 4-fold reduction of the RNA elongation product. These activity results are consistent with the notion that nsp8Δ76 can compete with full length nsp8 in the assembly of elongation-competent RdRp complex built from nsp12, nsp7, and nsp8.

### Use of AlphaFold-based in silico competition assay to search of nsp8 variants that confer stable nsp8-nsp12 interactions

Our function activity results are consistent with the idea that nsp8Δ76 can displace full length nsp8 in the RdRp complex. To optimize the interactions between nsp8 Δ76 to nsp12, we searched for single amino-acid mutations of nsp8 in the region (aa 104 – 131), a region in the C-terminal domain that confers protein-protein interactions with nsp12 in the reported structure [10]. To do so, we utilized AlphaFold (AF) ensemble competition assay for pairwise comparison between nsp8 single residue mutations and nsp8 wild type in binding to nsp12. Although AF does not directly predict the free energy associated with a protein-protein complex formation, it can provide relative comparison of binding interactions by placing the stronger binder in the binding site in the majority of the ensemble. (**Fig. 3A**). Each nsp8 amino acid residue selected for in silico mutational study was replaced by a representative subset of 14 amino acids (R, E, S, Y, N, G, I, K, H, D, C, M, F, and W) to span diverse side-chain characteristics; including positive and negative charges, hydrophobicity, steric bulkiness, and aromaticity. For each pairwise comparison of nsp8 single residue variation against parental nsp8, 100 models were generated. The number of models with mutant nsp8 bound to nsp12 for each mutant is summarized using a heat-map.

**Figure 3.**
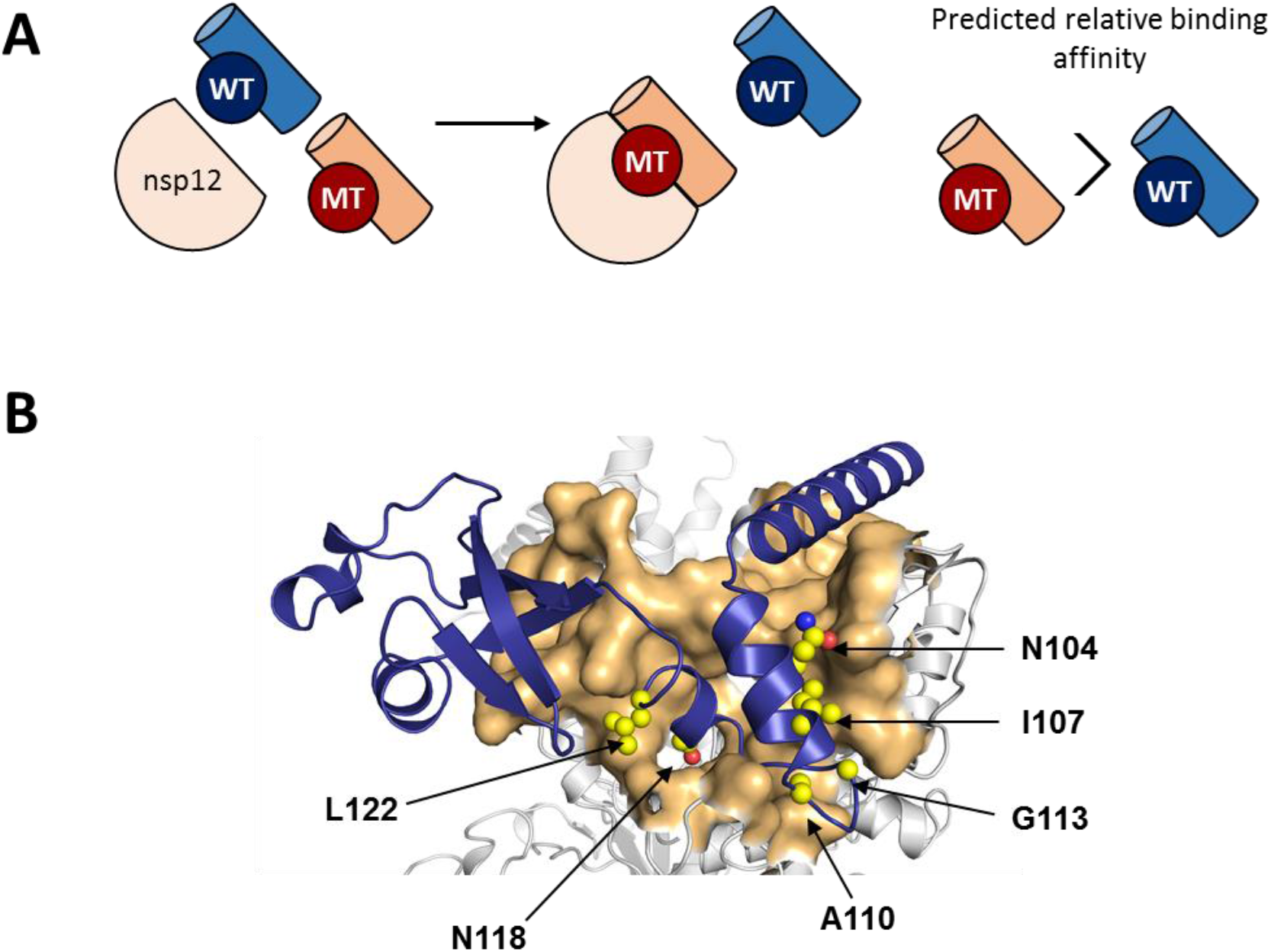
AlphaFold competitive binding assay. **(A)** Schematic representation of the AlphaFold ensemble competition assay design; WT – nsp8 wild type, MT – nsp8 mutant. **(B)** Detail of nsp12-nsp8 interface (PDB 6YYT). The hot-spot nsp8 residues (WT displaced by MT in more than 80 models out of 100) are shown as spheres.

The AF prediction identified several nsp8 mutations that were able to displace the parental protein in binding to nsp12 *in silico*. The single amino acid nsp8 mutants that displaced the wild type nsp8 include hotspots at N104, I107, A110, G113, N118, L122 with their locations at the interface with nsp12 annotated in **Fig. 3B**.

### Experimental evaluation of inhibitory behavior of the hotspot nsp8Δ76 mutants

The AlphaFold ensemble competition screens identified several nsp8 mutants were able to displace wild-type nsp8 in binding to nsp12. To test whether or not they would behave so in reality, we created the corresponding hotspot single amino acid mutants for nsp8Δ76 and measured their inhibitory potency using the *in vitro* RNA extension assay.

To evaluate whether or not these mutants can better inhibit RdRp RNA extension activity than wild type, we created hotspot single amino acid mutants for nsp8Δ76 at N104, I107, A110, G113, N118, and L122, and tested the inhibitory activity using the *in vitro* RNA extension assay using the RNA hairpin substrate. The RdRp RNA extension activity inhibition was measured by the amount of elongated RNA product by nsp12, nsp7, and nsp8 in the presence of the mutants (**Fig. 4**). The tested concentrations for the mutants were 6 μM, 12 μM, and 24 μM that corresponds to 1, 2, and 4-fold of wild type full-length nsp8.

**Figure 4.**
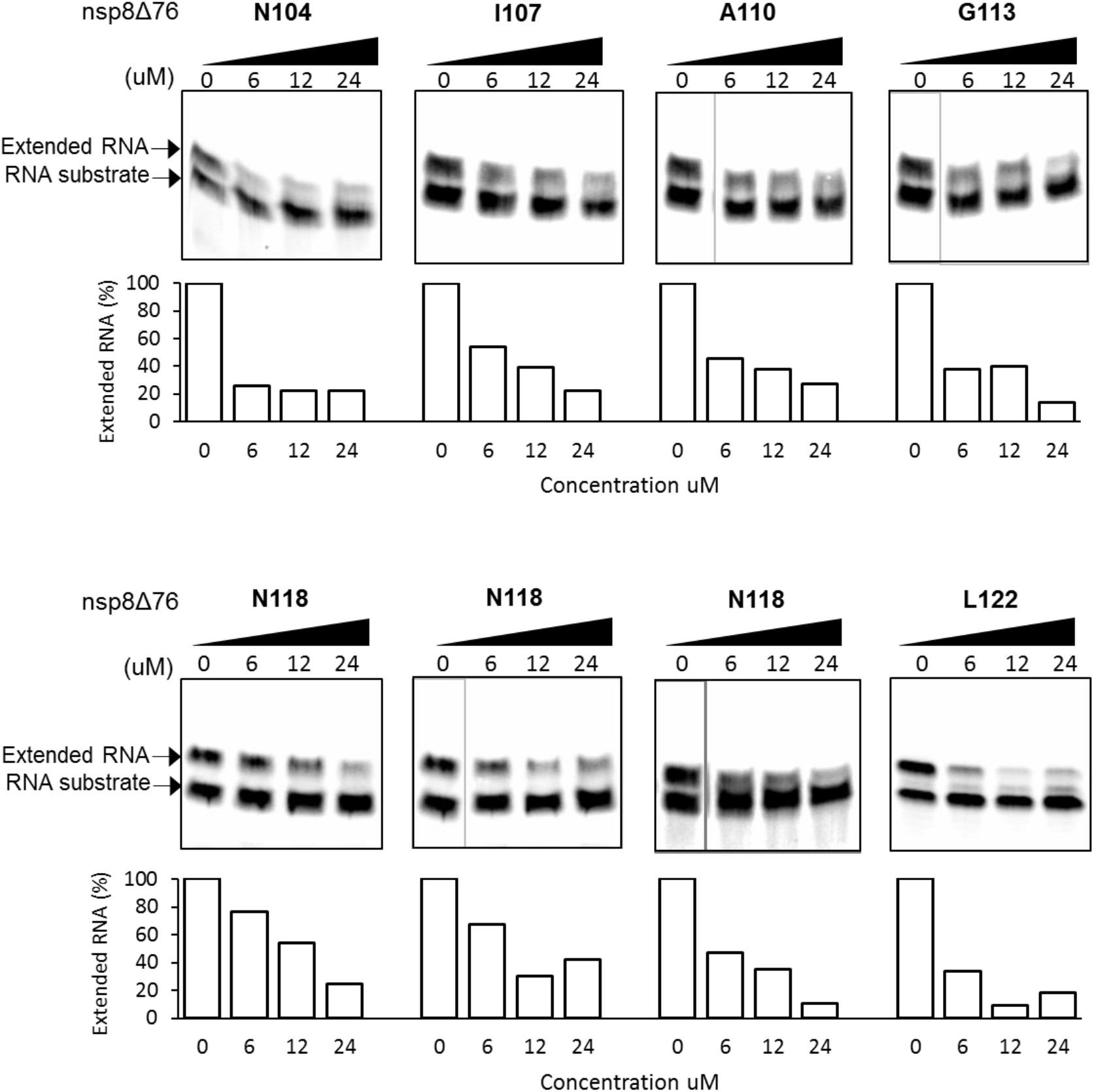
RNA elongation inhibition by nsp8Δ76 mutants. Up: RNA elongation reaction in a presence of nsp8Δ76 mutants. Bellow: Quantification of the fraction of extended RNA normalized to RdRp control reaction (without nsp8Δ76).

Most of the hotspot nsp8Δ76 mutants exhibit quantitative inhibitory behavior for RNA elongation across 6 μM, 12 μM, and 24 μM. Interestingly, a mutant at N104 that shows a modest score shows saturating inhibitory behavior across 6 μM, 12 μM, and 24 μM.

## Discussion

Pathogenic human coronaviruses have caused major outbreaks, such as recent severe acute respiratory syndrome coronavirus-2 that caused the recent COVID-19 pandemic [1]. Coronaviruses are positive-sense single-stranded viruses that rely on viral RdRp to replicate and transcribe viral RNA genome. Compared to other RNA viruses, the Coronavirus RdRp replicating system is a multiprotein complex, composed of nsp12, nsp7, and nsp8 subunits. The nsp8 subunit is critical for efficient RdRp processivity [3]. While we were preparing this manuscript, the conserved nsp12-nsp8 interface has been confirmed as a promising target for developing peptide inhibitors against RdRp as antiviral therapy [15]. In our study, we based on a structural rationale to target this nsp12-nsp8 interface, but further explore variants of nsp8-based protein fragment that can effectively compete out the wide type nsp8, thus disrupting the assembly of functional RdRp complex to suppress the RdRp RNA elongation activity.

### Nsp8Δ76 is able to inhibit the RdRp activity

In our study, we used in vitro reconstituted RdRp complex from bacterially expressed nsp12, nsp7, and nsp8. This reconstituted complex shows an RNA extension activity on minimal RNA hairpin substrate (**Figure 1**). The nsp8 N-terminal extensions are critical for RdRp complex stability and the reported structure of a replicating complex suggested that deletion of the N-terminal helix region may reduce the RNA elongation activity [10]. We thus created a nsp8 fragment with deletion of an N-terminal α-helix (Δ1-75), nsp8Δ76. This nsp8 fragment still binds to nsp12 (data not shown) but does not support RNA elongation activity (**Figure 2B**).

### Experimental evaluation of hotspot inhibitory nsp8Δ76 mutants predicted by AlphaFold competitive binding assay eliminates computational artifact

Using AlphaFold competition binding assays, we identified several nsp8 mutants with potentially better nsp12 binding than nsp8 wild type (**Figure 3**). To date, computational approaches are invaluable in facilitating effective search of protein mutants, in particularly useful for the design of peptide inhibitors [20,21]. The peptide binder can be optimized for higher affinity and increased inhibitory efficiency through enhancing interactions between peptide and protein through replacing the amino acids of peptide at the interface for optimizing protein-protein interactions. Using an AlphaFold-based computational approach recently proposed [17,18]. These mutations included hot-spot residues N104, I107, A110, G113, N118, and L122, consistent with the nsp12-nsp8 interface region identified through quantum mechanical calculations [12].

We further subjected those nsp8Δ76 mutants to *in vitro* RNA elongation assay (**Figure 4**). Among the selected nsp8Δ76 mutants, most of the mutants showed dose dependent reduction of RdRp elongation activity as wild type nsp8Δ76 except that at 104 conferring saturating inhibitory activity across different doses. In conclusion, our study demonstrated that a combined experimental and computational framework for targeting critical protein-protein interactions for inhibiting SARS-CoV-2 RdRp replication activity.

## Materials and Methods

### Bacterial expression and purification of recombinant protein nsp12

The codon optimized cDNA coding for nsp12 (GeneScript) was cloned into modified pETDuet-1 vector for expressing His6-SUMO-nsp12-strep-tag. The plasmid was transformed into *E. coli* Rosetta 2 (DE3) cells (ECOSTM) and grown overnight on LB-agar plates containing 100 μg/mL ampicillin (AMP) and 25 μg/mL chloramphenicol (CAM) antibiotics. Single colonies were used to inoculate 20 mL of LB/AMP/CAM medium and grown overnight at 37°C. This overnight culture was used to inoculate 2 L of LB/AMP/CAM medium and grown at 37°C until the optical density at 600 nm (OD@600) reached 0.6. The protein expression was induced by addition of IPTG to a final concentration of 0.1 mM and then incubated at 16°C overnight (16 hrs). Cells were harvested by centrifugation at 5000 rpm for 25 min at 4°C. Cell pellets were re-suspended in lysis buffer (20 mM Tris, pH 8.0, 500 mM NaCl, 5% (v/v) glycerol, 0.1 mM EDTA, 5 mM β-mercaptoethanol (β-ME), 1x complete Protease Inhibitor (RIP, Roche), and lysed using a high-pressure homogenizer. The lysate was cleared by centrifugation at 150 000 g, 30 min, at 4°C and incubated with Ni Sepharose Fast Flow (Cytiva), washed, and eluted with Nickel elution buffer (20 mM Tris, pH 8.0, 300 mM NaCl, 5% (v/v) glycerol, 500 mM imidazole, 5 mM β-ME). The eluted nsp12 was then incubated with Strep-tactin X resin (Cytiva), and the N-terminal His6-SUMO-tag was cleaved using Ulp1 SUMO protease and washed to remove the SUMO protease with wash buffer (20 mM Tris, pH 8.0, 300 mM NaCl, 10% (v/v) glycerol, 5mM MgCl2, 2 mM DTT). The nsp12 was eluted using SA elution buffer (20 mM Tris, pH 8.0, 300 mM NaCl, 5% (v/v) glycerol, 5mM MgCl_2_, 50 mM biotin, 5 mM β-ME) and loaded onto Superdex 200 Increase 10/30 GL equilibrated with gel filtration buffer (20 mM Tris, pH 8.0, 300 mM NaCl, 5% (v/v) glycerol, 5 mM MgCl_2_, 2 mM DTT). Purified nsp12 was concentrated using centrifuge concentrator with 30 kDa MWCO (Millipore) in storage buffer (20 mM Tris, pH 8.0, 300 mM NaCl, 10% (v/v) glycerol, 5 mM MgCl2, 2 mM DTT), aliquoted, and stored in −80°C.

### Bacterial expression and purification of recombinant protein of nsp7 and nsp8

The pETDuet-1 codon optimized (GeneScript) plasmids expressing His6-SUMO-nsp7 and His6-SUMO-nsp8 were transformed into *E. coli* BL21 (ECOSTM) grown in LB medium supplemented with 100 μg/mL AMP. Single colonies were used to inoculate 20 mL of LB/AMP medium and grown overnight at 37°C. This overnight culture was used to inoculate 2 L of LB/AMP medium and grown at 37°C until OD@600 reached 0.5. The protein expression was induced by addition of IPTG to a final concentration of 0.1 mM and then incubated at 16°C overnight. Cells were harvested by centrifugation at 5000 rpm for 25 min at 4°C. Cell pellets were re-suspended in lysis buffer (20 mM Tris, pH 8.0, 300 mM NaCl, 5% (v/v) glycerol, 20 mM imidazole, 5 mM β-mercaptoethanol (β-ME), 1x complete Protease Inhibitor (RIP, Roche), and lysed using high-pressure homogenizer. The lysate was cleared by centrifugation at 150 000 g, 30 min, at 4°C and incubated with Ni Sepharose Fast Flow (Cytiva), washed, and eluted with Nickel elution buffer (20 mM Tris, pH 8.0, 300 mM NaCl, 5% (v/v) glycerol, 500 mM imidazole, 5 mM β-ME). The N-terminal His6-SUMO-tag was cleaved using Ulp1 SUMO protease and cleaved N-terminal His6-SUMO-tag and SUMO protease was removed by incubation with Ni Sepharose resin. The nsp7 and nsp8 in the non-binding fraction was concentrated using centrifuge concentrator with 3 kDa MWCO (Millipore) and put into storage buffer (20 mM HEPES, pH 7.5, 150 mM NaCl, 5% (v/v) glycerol, 2 mM DTT), aliquoted, and stored in −80°C.

### nsp8Δ76 and single amino acid nsp8Δ76 variants

To create N-terminally truncated nsp8Δ76 construct, nsp8 sequence spanning amino acid (a.a.) 76-198 was amplified from pETDuet-1-nsp8 plasmid using PCR reaction. The amplified DNA fragment was cloned into a modified pETDuet plasmid with N-terminal His6-SUMO-tag. The new construct was verified by DNA sequencing. The nsp8Δ76 proteins was expressed in *E. coli* BL21 (DE3) (ECOS) and purified using the aforementioned protocol used for purification of full-length nsp8. nsp8Δ76 was concentrated and exchanged into storage buffer (20 mM Tris, pH 8.0, 150 mM NaCl, 5% (v/v) glycerol, 2 mM DTT) using centrifuge concentrator with 3 kDa MWCO (Millipore), aliquoted, and stored in −80°C.

Single amino acid mutations in nsp8Δ76 were constructed by PCR reaction using PCR primers coding targeted amino acid mutation and the DNA sequence of nsp8Δ76 mutant was confirmed by sequencing. The expression and purification of nsp8Δ76 mutants followed the aforementioned purification protocol used for purifying the wide type nsp8Δ76.

### RNA elongation assay

The minimal RNA scaffold used for *in vitro* RNA elongation reaction was purchased from Integrated DNA Technologies. The hairpin RNA substrate connecting the template RNA to the RNA primer through a tetraloop with the sequence 5’-Cy3-UUUUCAUGCUACGCGUAGUUUUCUACGCG-3’ was designed according to that in a previous work of RNA elongation assay [10]. RNA was annealed by heating to 75 °C and gradually cooling to 4 °C in the buffer containing 10 mM Tris, pH 8.0, 25 mM NaCl, 1 mM EDTA.

The RdRp complex was reconstituted by incubating nsp12 (1.0 μM), nsp7 (3.0 μM), and nsp8 (6.0 μM) at 4°C overnight. For RNA elongation reaction, the RdRp complex was incubated with RNA scaffold (1 μM) for 30 min at 30°C and the RNA extension was initiated by addition of NTPs (1 mM ATP, CTP, GTP, and UTP each) in 20 mM Tris, pH 8.0, 110 mM NaCl, 5% (v/v) glycerol, 10 mM MgCl_2_, and 1 mM DTT. Reactions were quenched by addition of 2x stop buffer (7 M urea, 50 mM EDTA, pH 8.0, 1x TBE buffer). Samples were digested with proteinase K (New England Biolabs). The reaction was loaded onto a 15% TBE-urea polyacrylamide gel (Thermo-Fisher), run at 150V for 90 min, and RNA products were visualized by iBright FL1500 Imaging System.

For RNA elongation inhibition reactions, the inhibitory nsp8 fragment was added to the RdRp-nsp7-nsp8 RNA complex and incubated for additional 30 min at 30°C prior to addition of NTPs to initiate the RNA elongation reaction.

### AlphaFold Computational Competition Assay

Modeling was performed on a local install of the publicly available AlphaFold 3 (version 2.0). For each specific mutation, 100 independent models were generated to assess the occupancy of the primary binding site on nsp12. The “binding success rate” was defined as the frequency with which the mutated nsp8 successfully occupied the target interface on nsp12 in the presence of a wild type nsp8 as a competitor. To cover a broad spectrum of physicochemical properties, we selected a representative subset of 14 amino acids (R, E, S, Y, N, G, I, K, H, D, C, M, F, and W) for mutational scanning. This subset was strategically selected to span diverse side-chain characteristics—including positive and negative charges, hydrophobicity, steric bulkiness, and aromaticity—ensuring that the competitive landscape reflects the fundamental chemical determinants of the nsp12-nsp8 interface. The results were combined and visualized in a heat-map, where each cell represents the binding occupancy count (ranging from 0 to 100) for a specific substitution at a given residue position.

